# A Mathematical Model of Persister Cell Plasticity and Its Impact on Adaptive Cancer Therapy

**DOI:** 10.1101/2025.06.26.661628

**Authors:** Adithya Mathews, David Basanta

## Abstract

The emergence of resistance to therapy remains a significant obstacle to successful treatment in cancer that is driven by somatic evolution. Adaptive therapies represent a novel approach to manage the emergence of resistance, by leveraging the competition between different cell phenotypes to control tumor burden rather than aiming for complete eradication. However, the emergence of phenotypica **l**y plastic persister cells, exhibiting transient epigenetic resistance, poses a significant challenge to the efficacy of these approaches, and their specific impact is often overlooked in preclinical and mathematical models. This study investigates the role of persisters within adaptive therapy using a spatial agent-based model simulating sensitive, persistent, and genetically resistant cell populations. Our simulations reveal that persisters critically undermine treatment efficacy, significantly reducing progression-free survival (PFS) by approximately 40% (average 207 vs. 344 days in simulations) as they provide a reservoir for acquiring genetic resistance. Furthermore, tumors with higher levels of epigenetic resistance (more robust persisters) showed accelerated evolution towards resistance dominance phenotypes. The model showed treatment was most effective when sensitive cells initially dominated the tumor microenvironment. These findings highlight that persister dynamics are crucial determinants of adaptive therapy outcomes, suggesting that future strategies must account for epigenetic resistance, potentially informing approaches to assess tumor composition and sensitivity to better tailor treatments and improve patient outcomes.

## 1. Introduction

A central challenge in oncology is the evolution of treatment resistance, which limits the long-term efficacy of many cancer therapies. Even initially successful treatments can eventually fail as tumor cell populations adapt and evade therapeutic pressure. Central to this challenge is the fact that cancers display a high degree of intra tumor heterogeneity (ITH) which makes it hard for treatments to completely eradicate a critical number of tumor cells which would be required to ensure the tumor does not come back. This ITH often manifests in terms of genetic mutations, but epigenetic alterations (such as methylation) are also possible. Further complicating this picture is the role of environmentally mediated drug resistance (EMDR) which allows for otherwise treatment sensitive tumor cells to survive the onslaught of treatment.

This complexity makes the study of the emergence of resistance to treatment in cancer amenable to mathematical modeling approaches [1], [2], [3]. This is especially true when it integrates clinical and pre-clinical data to ground these models in reality [4], [5]. Integration of experimental data in mathematical models allows for the generation of novel hypotheses that can be tested in a lab [1], [2].

On such hypothesis derived from mathematical modeling led to the concept of Adaptive Therapies (AT), a treatment modality in which tailoring the treatment approaches and dosages according to tumor growth, is an accessible tool to handle the possible emergence of resistance [6], [7], [8]. This approach holds promise in stabilizing treatment-resistant cell populations, potentially leading to an extension in patient survival but assumes that intrinsic resistance results from genetic mutations, which could provide resistance but also represent in some cases a fitness cost.

Cancer cells have been shown to exhibit phenotypic plasticity, allowing them to quickly change phenotypic characteristics in response to environmental stimuli, without genotypic changes. This adaptability greatly contributes to the ability of persister cells to survive treatment [8], [9]. This phenotypic plasticity represents a critical factor in the complex landscape of somatic evolution within tumors [10] and highlights the challenges posed by intertumoral heterogeneity in the context of cancer therapeutics and emphasizes the need for sophisticated treatment strategies that account for the dynamic nature of tumor subpopulations. The processes behind persisters behavior are not fully elucidated and present a challenge to effective therapy design [8].

While the full extent of the interplay between persister cells and treatment is unknown, there is evidence that the survival of persistent cells during treatment could allow for the acquisition of genetic mutations leading to genetic drug resistance, further compromising treatment efficacy [7], [8], [9]. Alternatively, some studies have shown evidence for the possibility that some cancer patients could regain drug sensitivity following a treatment hiatus—commonly referred to as a “drug holiday”—suggests an epigenetic basis for transient drug tolerance [7]. The detection of a distinct chromatin state in drug-tolerant cancer cells and consequent hypersensitivity to HDAC inhibitors potentially yields a therapeutic opportunity to prevent the development of stable drug resistance [7]. The creation of persistence would be a precursor to drug resistance; therefore, an understanding of this step can decrease the chance of the tumor developing resistance [9].

Addressing the challenge of persisters in cancer treatment requires novel approaches. Strategies targeting the mechanisms that mediate the transition to a persister state during drug treatment may help prevent the emergence of drug resistance. Additionally, understanding the unique metabolic and regulatory characteristics of persisters could reveal exploitable therapeutic vulnerabilities [9] This study aims to focus on the unique role of persisters in contributing to tumor resilience and potential treatment failure in the case of adaptive therapies. The reasoning puts persisters at the forefront of therapeutic resistance, emphasizing their ability to direct adaptive treatment strategies that seek to reverse tumor evolution dynamics.

## 2. An agent-based model of epigenetic resistance in cancer

The model uses a 100×100 grid (‘lattice’) where 9 sensitive cells are centrally placed at the start of each simulation. The model considers three different cell subpopulations—sensitive, resistant, and persistent—differentiated from each other by their response to treatment. Simulations were run for 350 timesteps, allowing tumor growth and treatment dynamics to unfold. Cells undergo proliferation or apoptosis stochastically, completing one cycle per timestep reaching a carrying capacity of approximately 9000 cells (see supplementary figure).

Tumor cell proliferation requires available space, while apoptosis frees it. Sensitive cells can mutate into persistent or resistant cells, although the rate of mutation from sensitive to resistant cells is significantly lower than that of sensitive to persistent cells. Persistent cells act as an intermediate pool for resistance development, providing an easier path for resistant cells to emerge [8], [9]. In this model, the mechanisms through which persistent cells develop resistance to treatment are not considered as there are diverse number of mechanisms through which epigenetic resistance can be expressed. To ensure these results are applicable to different types of cancer, we assume that all cells are equally predisposed to persistence. Sensitive cells exhibit baseline proliferation and apoptosis rates, altered under treatment. Resistant cells maintain constant but slower proliferation due to fitness costs. Persistent cells display phenotypic plasticity, mirroring sensitive cells in untreated conditions but showing partial resistance under treatment.

Adaptive treatment adjusts based on tumor size, regulating tumor population between a start (*N*_*start*_) and end (*N*_*stop*_) population. Cell dynamics are visualized through time-sequenced images, with subpopulations represented by distinct colors (Blue=Sensitive cells, Green=Persistent cells, Orange=Resistant cells). Cell counts are recorded at each timestep, providing data for analyzing treatment and growth patterns.

The goal of this work is to qualitatively explore how different types of self -intrinsic resistance impact treatment and to help understand the dynamics that drive the emergence of treatment resistant populations in cancer in general. As a result, model parameters used are not derived from experimental data. The model also simplifies types of resistance into two broad categories: epigenetic resistance and genetic resistance, which does not account for the full complexity of resistance in tumor biology.

The proliferation rate for sensitive, resistant and persister cells are denoted by 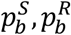 and 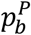 respectively. The apoptosis rates are represented by 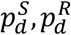 and 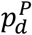. In general, the proliferation and apoptosis rate of cell types *C* ∈ { *S*, *R*, *P* } is symbolized by 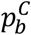 and 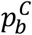. The transition rate or the mutation rate between subgroups where sensitive cells may mutate into resistant 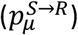 or persistent 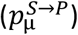 cells and persistent cells may mutate into resistant cells 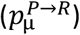. Treatment is modeled to be a time-dependent function that modulates the proliferation or apoptosis rate. The equations below indicate the effective probabilities of cell functions:

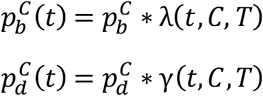

λ(*t*, *C*, *T)*, where *t* is the timestep, *T* as the type of treatment and *C* is the type of cell, represents the proliferation scaling factor and lies between and lies between 0 < λ < 1. γ(*t*, *C*, *T)* represents that apoptosis scaling factor and γ > 1. Detailed parameter space exploration was conducted to ensure that these outcomes are consistent across various parameter values, see supplementary analysis.

Sensitive subpopulation

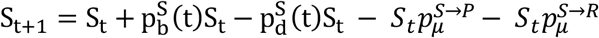

Resistant subpopulation

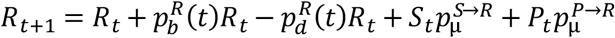

Persister subpopulation

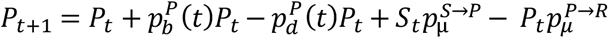

## 3. Results

### 3.1 Persisters Reduce the Likelihood of PFS

Persistent cells exhibit phenotypic plasticity, allowing them to survive drug treatment and contribute to tumorresilience [5], [8], [9], [11]. Unlike fully resistant cells, persisters can revert to a drug-sensitive state after treatment cessation, making them difficult to eradicate completely [7], [9]. This ability to survive initial treatment and potentially give rise to resistant populations significantly complicates cancer therapy strategies.

The presence of persisters in tumors creates a reservoir for the potential emergence of drug-resistant mutants. This phenomenon parallels observations in antibiotic-treated bacterial populations, where persisters play a crucial role in the development of antibiotic resistance.

The inclusion of persistent cells in tumor populations significantly reduces progression-free survival (PFS) in cancer treatment, as evidenced by the decrease in average PFS from 344 days to 207 days when persistent cells are present. This substantial reduction in PFS, as seen in Figure 3, underscores the critical role of persisters in cancer therapy challenges.

**Figure 1.**
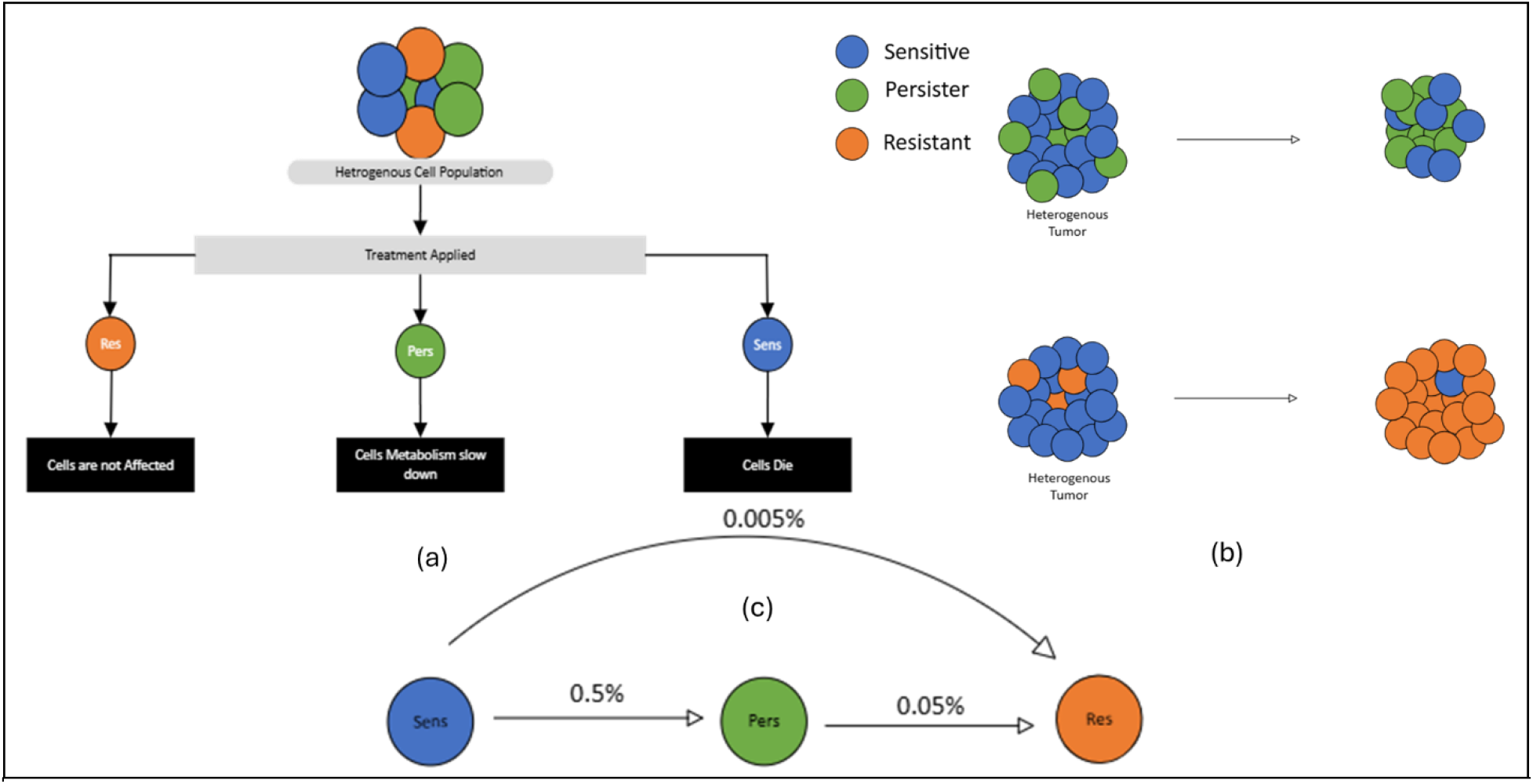
The multi-panel figure elucidates the model’s key features. (a) The left panel depicts a schematic representation illustrating the differential impact of the therapeutic intervention on distinct cellular subpopulations. (b) The middle plot compares tumor growth outcomes under treatment, based on different resistance subpopulations in the tumor and the expected tumor growth. (c) The middle plot displays the rate of mutation between the different subpopulations.

**Figure 2.**
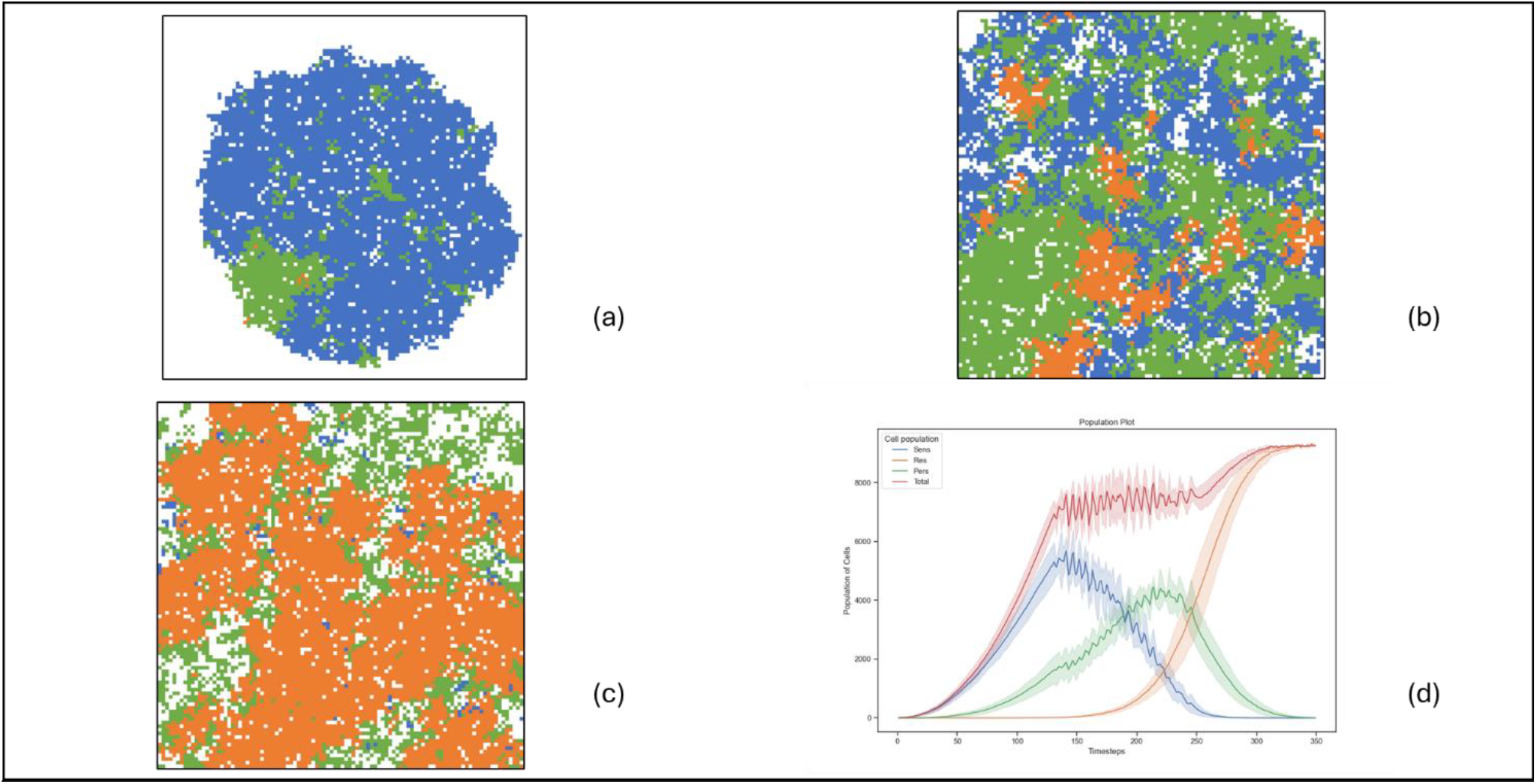
The graphical sequence illustrates the spatiotemporal evolution of tumor heterogeneity, while also highlighting the overall growth trajectory and subpopulation trends over time. These slides visually depict the tumor’s growth dynamics, with blue points representing sensitive cells, green points denoting persistent cells, and orange points indicating resistant cells, progressing sequentially from panel (a) to panel (c). Panel (d) illustrates the tumor’s growth through a plot, highlighting the population trends of each cellular subpopulation over time. These are the important parameters for the above simulation λ(*t*, *S*, *A) =* 0.4, λ(*t*, *P*, *A) =* 0.7, γ(*t*, *S*, *A) =* 1.75, γ(*t*, *P*, *A) =* 1.5 where *A* represents Adaptive therapy with *N*_*start*_ *=* 7500 and *N*_*stop*_*=* 7000. Additional parameter details are available in the supplementary material.

**Figure 3.**
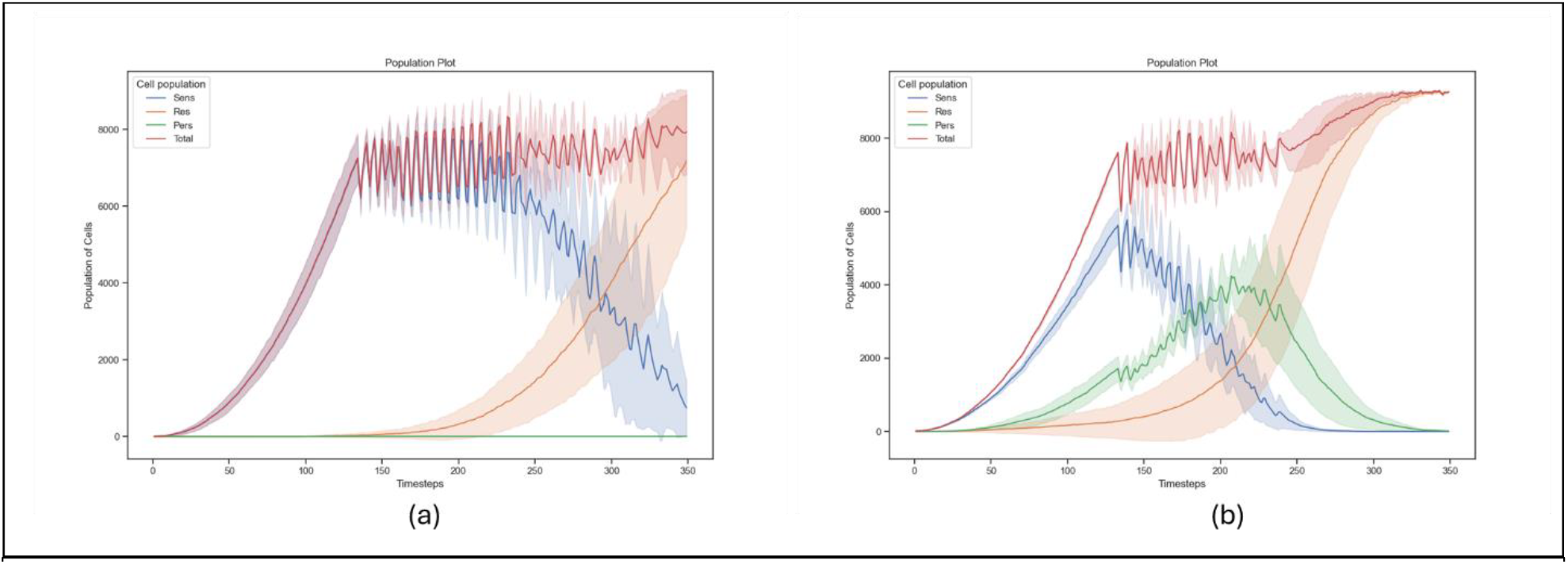
illustrates that the inclusion of a persistent cell population significantly reduces progression-free. (a) The left plot shows tumor growth with only sensitive and resistant populations, measuring progression-free survival (PFS) time based on RECIST criteria, with an average PFS of 344 days. (b) The right plot shows tumor growth with sensitive, persistent, and resistant populations, measuring progression based on RECIST, with an average PFS of 207 days.

Persistent cells indeed act as a reservoir for the potential emergence of drug-resistant mutants, facilitating an earlier onset of resistant phenotype dominance. Unlike fully resistant cells, persisters can revert to a drug-sensitive state after treatment cessation. This bidirectional phenotypic switching confers a survival advantage in both treatment-naive and treatment-exposed microenvironments, contributing to their role in tumor persistence and potential treatment failure.

### 3.2 Increased Epigenetic Resistance Correlates with Decreased PFS

In this model, to account for the dormancy of persistent subpopulation, which is achieved by a reduced sensitivity to treatment. To measure the efficacy of persisters correctly, we consider varying levels of sensitivity.

As persistent cell resistance increases, a significant spike in their population occurs. Conversely, when persisters exhibit higher treatment sensitivity to therapeutic interventions, there is decreased progression-free survival (PFS). Figure 4 (b) highlights the effect of epigenetic resistance level on the evolution of the tumor and subplot (a) given the population of persistent population based on the level of epigenetic resistance.

**Figure 4.**
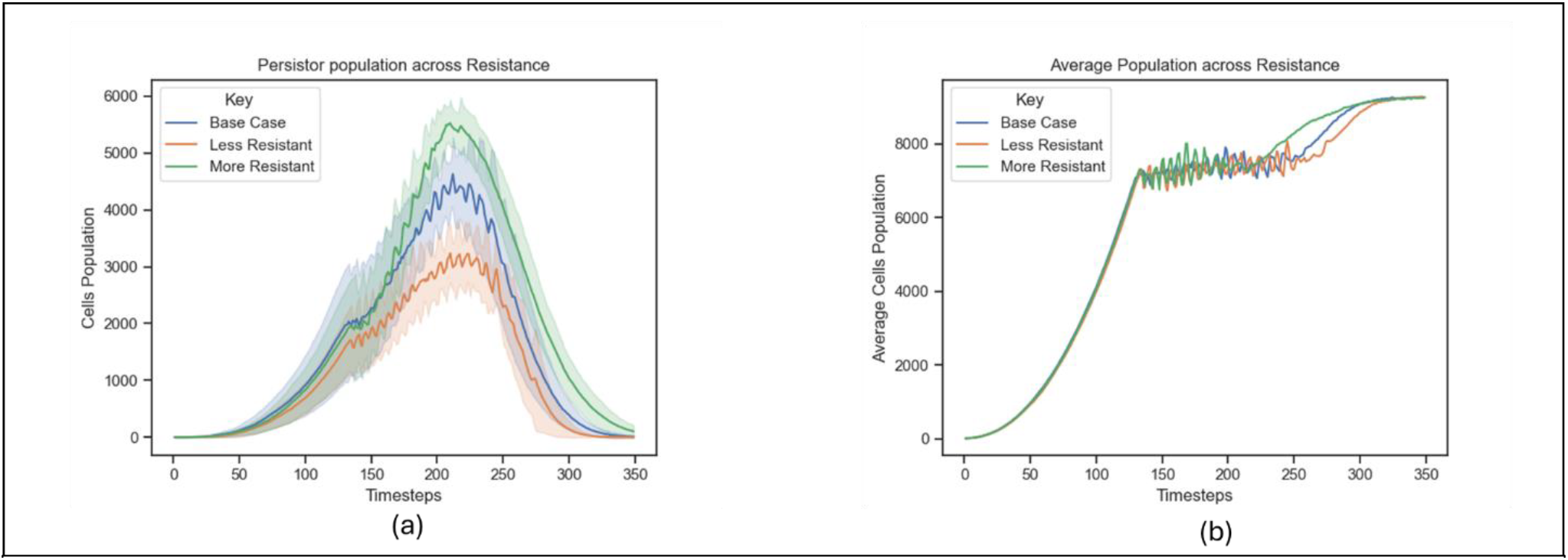
demonstrates that increasing resistance levels in persister cells lead to larger tumor populations and shorter PFS times, highlighting the impact of drug resistance on treatment efficacy. (a) The left plot compares varying resistance levels in persistor cells, showing an increase in population as resistance intensifies. (b) The right plot illustrates the average tumor population across different resistance levels, indicating shorter progression-free survival (PFS) with higher resistance.

The correlation found between the level of resistance in persisters and PFS is as anticipated to the treatment regime. The higher the capability of persistent cells for resisting apoptosis specific to treatment would cause the decline in the PFS, which establishes resistant subpopulation. The resistance phenotype of tumor occurs more widespread at a faster rate.

### 3.3 Experimental treatment to expose effect of epigenetic resistance on Drug Holidays

The experimental therapeutic module, as outlined below, is designed to identify the degree of heterogeneity of the tumor by varying treatment cycles and observing population dynamics of the tumor. It yields information regarding sensitive, persistent, and resistant subpopulations of cells within the tumor. The treatment process starts with the tumor when it is 5,000 cells and proceeds until the sensitive population is below 250 cells (5% of the initial population). This strategy effectively eliminates the majority of sensitive subpopulation. The subsequent drug holiday provides time for regrowth and re-sensitization of the tumor before re-initiation of treatment. Contrary to the initial hypothesis, the observed growth patterns deviated from expectations. According to the hypothesis, if persisters were present (epigenetic and genetic resistance), the tumor population would initially dip and then rise over time. Conversely, without persisters (genetic resistance), the population was expected to continuously increase after the second treatment. However, as seen in Figure 5 (a) and (b), the results showed a different pattern, suggesting that the underlying mechanisms of tumor heterogeneity and treatment response are more complex than initially theorized.

**Figure 5.**
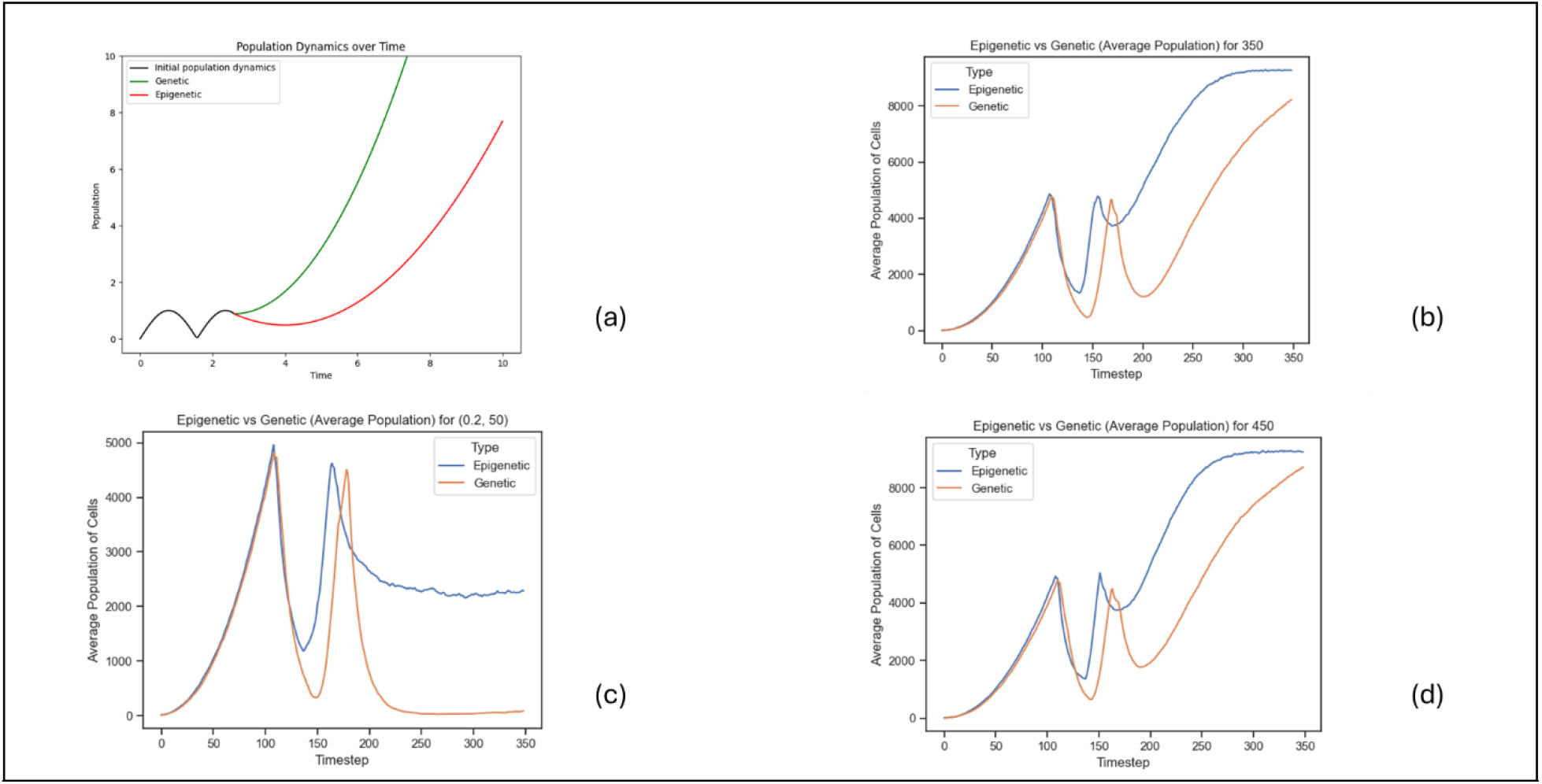
establishes setup of experimental treatment, expected treatment outcome along with resultant treatment outcomes. (a) The top-left plot provides a schematic representation of the anticipated tumor response upon application of the experimental treatment. (b) The top-right plot illustrates the projected outcomes following treatment administration. (c) The bottom-left plot demonstrates the scenario when the cost of resistance is high, while (d) the bottom-right plot depicts outcomes when the resistance cost is minimal.

To address the discrepancy between hypothesized and observed results, simulations were run across a range of two key parameters: length of drug holiday and resistance costs. The length of the drug holiday did not significantly impact the treatment outcome. The cost of resistance was a crucial parameter. It resulted in a suboptimal rate of proliferation of resistant cells to the optimal rate of proliferation of sensitive cells on treatment, resulting in flatlining of tumor growth as indicated by Figure 5 (c). Conversely, when the cost of resistance was lower, the resistant cell proliferation rate exceeded that of the sensitive subpopulation, resulting in continued tumor growth as seen in Figure 5 (d). To counterpoint the difference between postulated and attained results, simulations were carried out over a series of modifying two parameters: drug holiday length and resistance costs. The duration of the drug holiday was not so much to be blamed for the treatment out.

### 3.4 Adaptive Therapy Effective depended on Initial Tumor Sensitivity

The efficacy of adaptive therapy is intrinsically linked to the size of the tumor. The proportion of treatment-sensitive cells within the tumor microenvironment allows for greater control of the tumor growth and decay as well as tumor heterogeneity. The addition of epigenetic resistance into the equation adds complexity. In order to analyze whether a tumor is a suitable candidate for adaptive therapies, we can apply a small perturbation in order to access the level of sensitivity of the tumor.

Our analysis reveals a critical interplay between the resistance ratio (α, representing the fraction of resistance that is epigenetic) and the tumor’s initial sensitivity profile. Specifically, the impact on treatment response is most noteworthy in tumors with a lower percentage of sensitive cells. Conversely, as the proportion of sensitive cells approaches unity as indicated in Figure 6 (a), the rate of tumor reduction due to treatment converges regardless of resistance ratio. Figure 6 (b) illustrates the 95% confidence interval of the rate of tumor reduction across all values of alpha, which is significantly small at unity.

**Figure 6.**
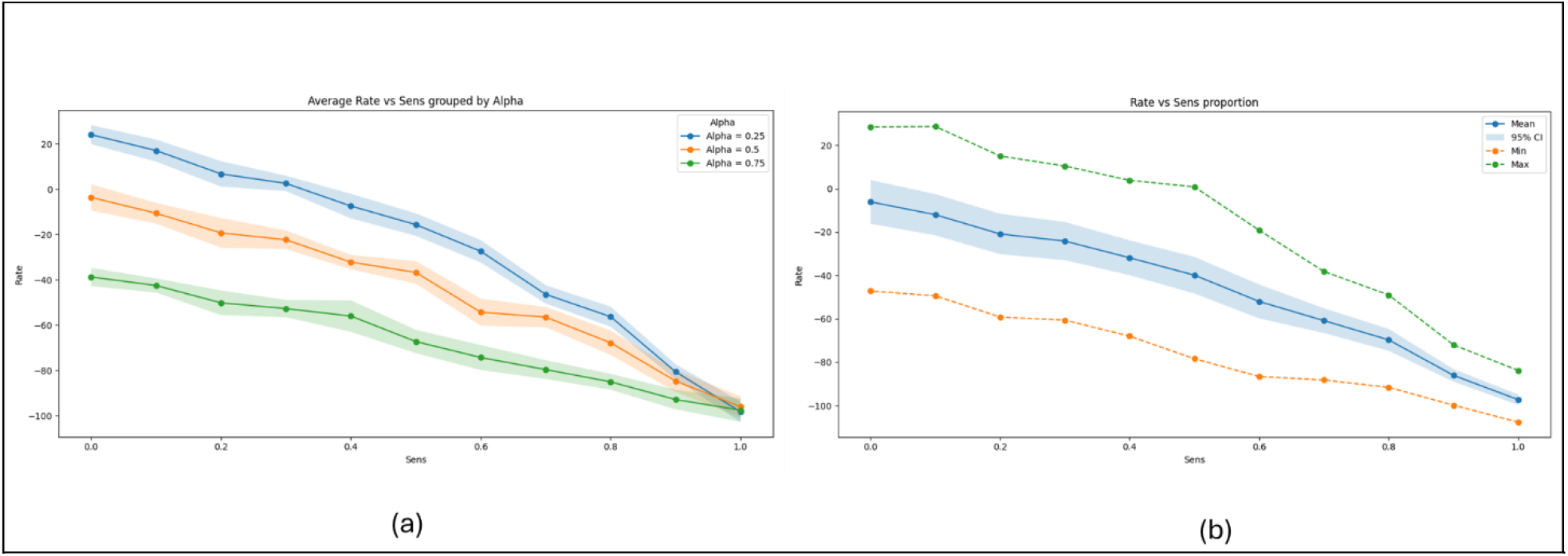
illustrates the influence of the sensitive cell proportion and resistance ratio (α) on tumor dynamics. (a) The left panel shows the average rate of tumor reduction versus the proportion of sensitive cells, grouped by different values of α, displaying the average rate converges as sensitivity increases. (b) The right panel displays the overall relationship between the rate of tumor reduction and the proportion of sensitive cells, highlighting the 95% confidence interval and the minimum/maximum reduction rates observed across simulations.

Hence if the perturbation of treatment leads to a rate of tumor decay that is within the ranges of treatment-sensitive dominant tumors, then we can affect epigenetic resistance is minimized and adaptive treatment is a viable option.

## 4. Discussion

Adaptive therapies minimize the emergence of fully treatment resistant tumors by leveraging the competitive dynamics between treatment-sensitive and resistance populations, not allowing tumors to become resistant phenotype dominant. However, existing mathematical models primarily focus on genetic resistance, failing to account for the role epigenetic resistance plays in tumor progression. Our study incorporates epigenetic resistance via the introduction of persistent cells into the model, addressing the growing body of evidence that persistent cells—characterized by phenotypic plasticity—play a significant role in treatment failure.

To investigate the impact of epigenetic resistance, we conducted simulations using an experimental therapeutic protocol. Aiming to treat a 2D tumor culture, this treatment is designed to isolate resistant populations in a tumor culture by diminishing treatment of sensitive populations to non-viable percentage of the population, preventing regrowth. Measuring these subpopulations separately, we can track the growth of each subpopulation [12], [13]. This protocol aimed at distinguishing tumors with and without epigenetic resistance by manipulating treatment cycles and observing tumor population dynamics. When persisters were present, tumor populations initially declined but later rebounded due to phenotypic switching and persistence, informing us about the potential of persistence to negatively impact tumor remission. It allows the tumor to transition from a sensitive dominant to resistance dominant state at a rapid pace.

Persistent cells act as a reservoir for potential resistance, significantly reducing progression-free survival (PFS). Unlike genetically resistant cells, persistent cells can revert to a drug-sensitive state after treatment cessation. This bidirectional phenotypic switching allows them to survive drug exposure and contributes to tumor resilience. Our results indicate that the presence of persistent cells reduces PFS substantially, highlighting their critical role in complicating cancer therapy strategies. By serving as a pool of resistance during treatment, they accelerate the dominance of resistant populations within the tumor. This parallels observations in antibiotic-treated bacterial populations, where persistent cells contribute to antibiotic resistance [9]. The phenotypic plasticity exhibited by these cells has variations across individual tumors and the degree of plastic behavior, i.e., the degree of epigenetic resistance plays a crucial role in the tumors ability to evade treatment. Our results demonstrate that the degree of epigenetic resistance significantly impacts treatment outcomes. Specifically, we observe that persistent behavior can be quantified by examining the transition rate from sensitive to persistent cells. These transitions not only allow tumors to escape treatment and drive resistance. By fitting experimental data to our model, we predict that the majority of persisters emerge upon exposure to targeted therapies. With new methods of estimating degree of resistance, it is worthwhile exploring quantification of the degree of epigenetic resistance, which might aid in personalized treatment protocols [13]. This finding aligns with current hypotheses regarding tumor heterogeneity and suggests that persistent cells play a pivotal role in adaptive therapy failure.

Given these findings, it is critical to determine whether a tumor is a suitable candidate for adaptive treatment. The effectiveness of adaptive therapy is closely tied to tumor size and sensitivity profiles. Larger tumors with a higher proportion of sensitive cells allow for greater control over the growth and decay dynamics. However, when epigenetic resistance is introduced into the model, this control diminishes due to the added complexity of phenotypic switching. Our analysis highlights a critical relationship between the resistance ratio (α) and the tumor’s initial sensitivity profile. Tumors with lower proportions of sensitive cells are more affected by α than those with higher proportions. As the proportion of sensitive cells approaches unity, treatment response becomes less dependent on α, suggesting that adaptive therapy remains viable when sensitivity dominates. To determine whether a tumor is suitable for adaptive therapy in practice, small perturbations can be applied to assess its sensitivity range. If these perturbations result in decay rates consistent with treatment-sensitive dominant tumors, it can be inferred that epigenetic resistance is minimized, and adaptive therapy is likely effective.

The formulated model presents the following important strengths. Firstly, it addresses spatial heterogeneity, a critical factor in accurately representing the intricate tumor microenvironment, allowing for a nuanced understanding of different regions within which the tumor may behave. Second, incorporating stochasticity into the model allows it to account for the underlying variability in biological systems by encapsulating the variety of possible results and better representing the dynamics of tumor behavior. A novel aspect of the model is the introduction of persistent phenotypic cells. This represents an advancement in modelling tumor heterogeneity, which could lead to new insights into major issues in cancer treatment such as tumor recurrence [5]. By incorporating epigenetic adaptation, the model has the potential to better predict long-term tumor behavior and guide future treatment protocols. Finally, the visualization capabilities offer a powerful tool to better comprehend tumor growth patterns and facilitate a deeper understanding of spatial and temporal dynamics. It is a communication tool enabling one to visualize the intricate behavior of the tumor microscopically, in an easily understandable manner, in a readily comprehendible fashion.

While the model provides valuable insights into tumor dynamics and treatment response, it is essential to acknowledge its limitations to ensure a comprehensive understanding of its applicability and interpretability. The most notable assumption is the simplification of biological process, as stated in the model description, including the simplification of condensation of types to resistance to two umbrella categories and assuming the impact of persistent treatment evasion mechanisms are irrelevant to the overall outcomes as persistent independently evolves different mechanisms to a singular treatment [8]. One of the primary limitations of this study is that we have not sought experimental or clinical validation of our results. Our goal is to provide a theoretical framework to be used to interpret the differential role of genetic and epigenetic resistance to treatment in a variety of cancer contexts. Recent work has been done to produce quantitative mathematical approaches that could capture more accurately some of these dynamics in specific cancer contexts [5]. Cell migration patterns have not been accounted for in the model framework. Cell migration plays a crucial role in tumor progression and metastasis, influencing treatment outcomes and overall disease prognosis. By neglecting this aspect, the model fails to capture the full spectrum of tumor behavior, potentially limiting its predictive capabilities and relevance to clinical scenarios. Furthermore, the simplification of RECIST criteria to a single condition overlooks the multifaceted nature of tumor response assessment. RECIST encompasses various parameters, including tumor size, nodal status, and presence of metastases, which collectively inform treatment decisions and prognosis. By reducing RECIST to a singular criterion, although necessary due to nature of the model, the model may oversimplify treatment response evaluation, potentially obscuring nuances that could impact treatment efficacy and patient outcomes. The assumption that all cells have the same distribution of epigenetic response is not reflected in clinical observations, as epigenetic plasticity is highly dependent on both the type of cancer and the specific treatment administered [14]. This necessitates the identification of principles that are held true across various treatment modalities and diverse tumor types.

Some potential avenues for exploration to address the limitations of the model are the incorporation of data-driven models into tumor progression so that it is based on real world data [3]. This approach has the added advantage of potential personalization of treatment strategies. Alternatively, using experimental data to validate the model’s projected tumor growth will ensure robustness [5].

Another avenue of further research is to conduct clinical trials to validate the method of assessing tumor suitability for adaptive therapy. Application of small perturbations to tumors and assess their sensitivity range to note whether the assertion made in the model holds true in the clinical landscape. These trials confirm whether tumors with minimal epigenetic resistance are indeed more suitable for adaptive therapy. We can quantify resistance for persister like HSLCI [13]. It quantifies resistance by measuring changes in cell biomass over time in response to drug treatment. It uses optical cell biomass measurements to track individual cell growth in real-time, and monitors thousands of individual cells or organoids simultaneously. It analyses growth patterns to identify cells that are persistently sensitive to specific treatments [12]. Exploration of these quantifications techniques informs personalized treatment protocols and improve adaptive therapy strategies.

In conclusion, our findings emphasize the need for a more nuanced approach to cancer treatment that accounts for the complex dynamics of persistent cell populations. By better understanding and targeting these cells, we may be able to develop more effective therapeutic strategies that extend progression-free survival and improve patient outcomes.

## Code Availability

The computational model code is available in an online repository at https://github.com/A-Binoy/Cancer-sim-python.

## Acknowledgements

We are grateful to Moffitt Cancer Center and the Department of Integrated Mathematical Oncology for supporting this research. We would like to thank Dr. Thomas Bieske and Dr. Catherine Beneteau (in the department of Mathematics and Statistics at the University of South Florida) for their advice and guidance.

## Supplementary Analysis

### Exploring Parameter Space

**Supplementary Figure 1.**
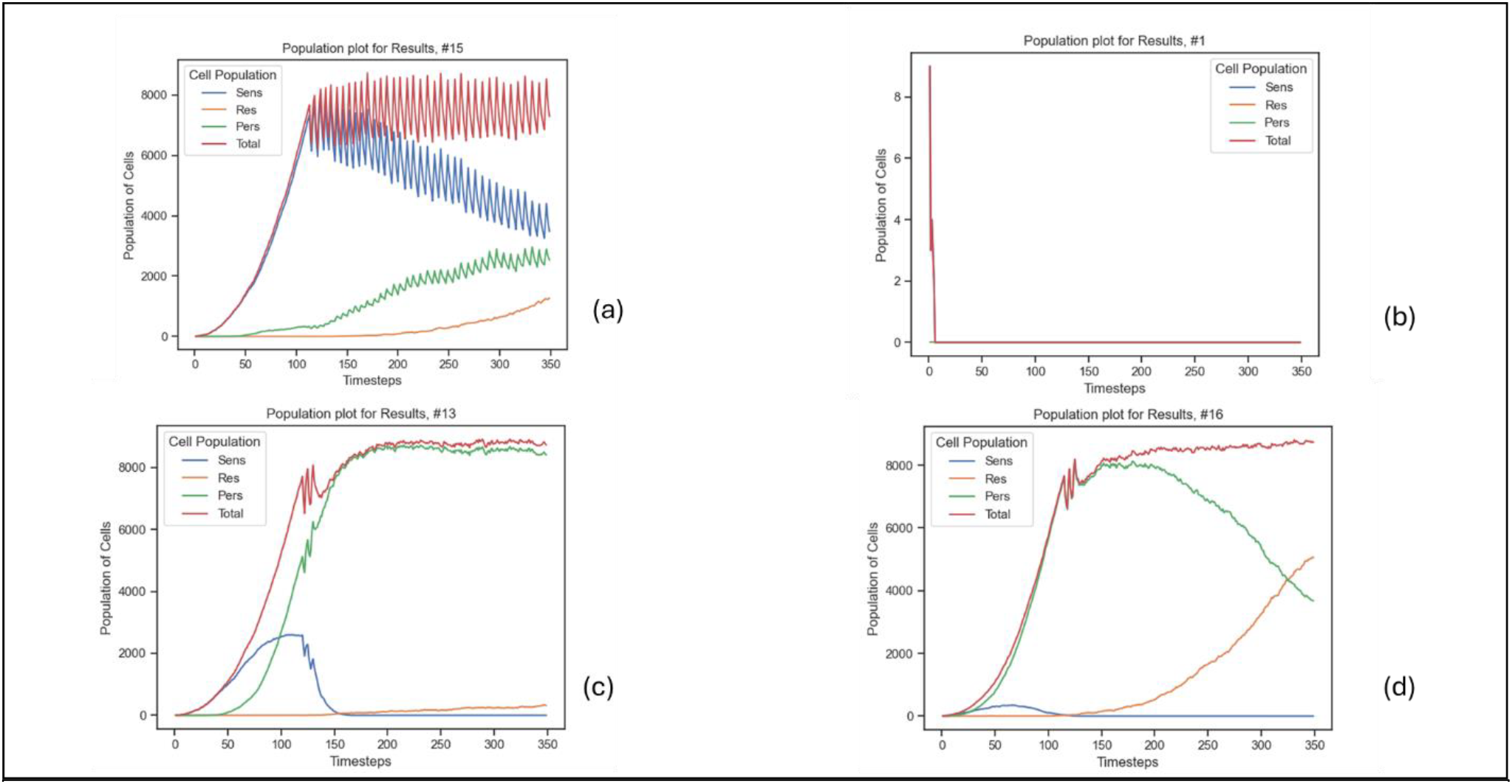
This figure illustrates tumor growth simulations under random parameter. (a) demonstrates oscillatory growth dominated by sensitive cells, while (b) shows tumor remission. (c) Sensitive cells initially increase but are eventually depleted, allowing persistent and resistant cells to dominate, leading to a stabilized total population. (d) depicts a tumor in which persistent cells dominate the tumor’s initial growth with a gradual shift toward resistant cell dominance.

**Supplementary Figure 2.**
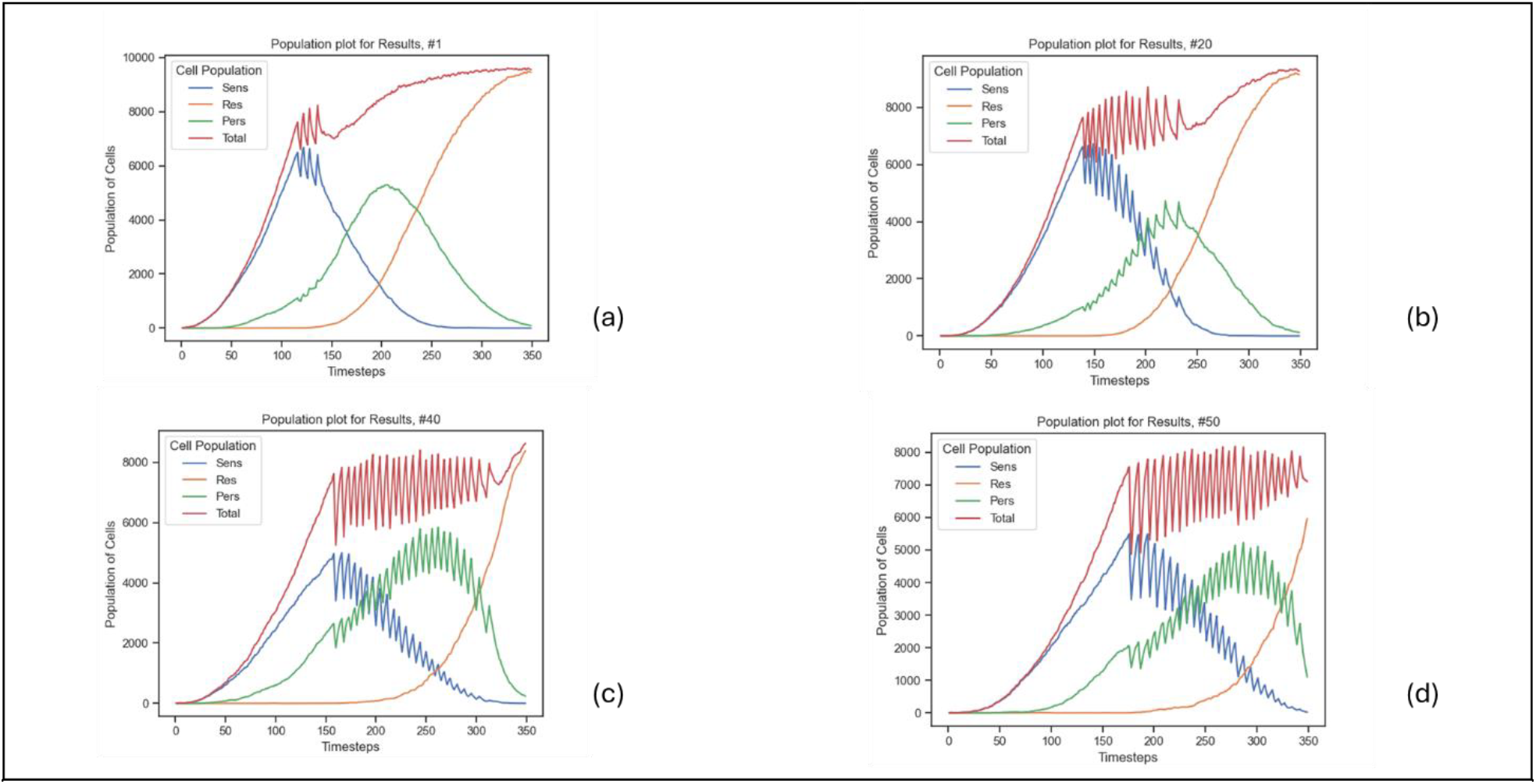
display symmetric shifts of ideal parameter values in a parameter range. From (a) to (d) the parameter values increase increasing the oscillations. The sequential panels, (a) through (d), demonstrate a progressive increase in parameter values, in tandem amplifying the magnitude and frequency of oscillations in the system under investigation.

The findings illustrated in Figures 1 and 2 underline that the primary determinants of the observed tumor growth dynamics are not individual parameters values. The dynamics hold consistent across all a range of parameter values. This allows us to make generalizations that are applicable to a wide range of tumors. Figure 2 varies parameter values within a range of biologically and clinically relevant ranges with symmetric shifts. This indicates that the models dynamics are not dependent on fine -tuning individual parameters. The observation suggests that the model captures intrinsic population dynamics rather than an artifact of specific parameter settings. Such robustness increases confidence that the model’s predictions are generalizable.

Here is a list of parameters that have been derivedand used in the model, baseline proliferation rates are 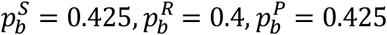. The baseline apoptosis rates are 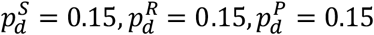.

The mutations rates are 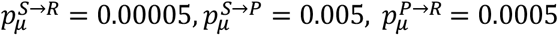.

### Negligent impact of Plastic Response Delay

**Supplementary Figure 3.**
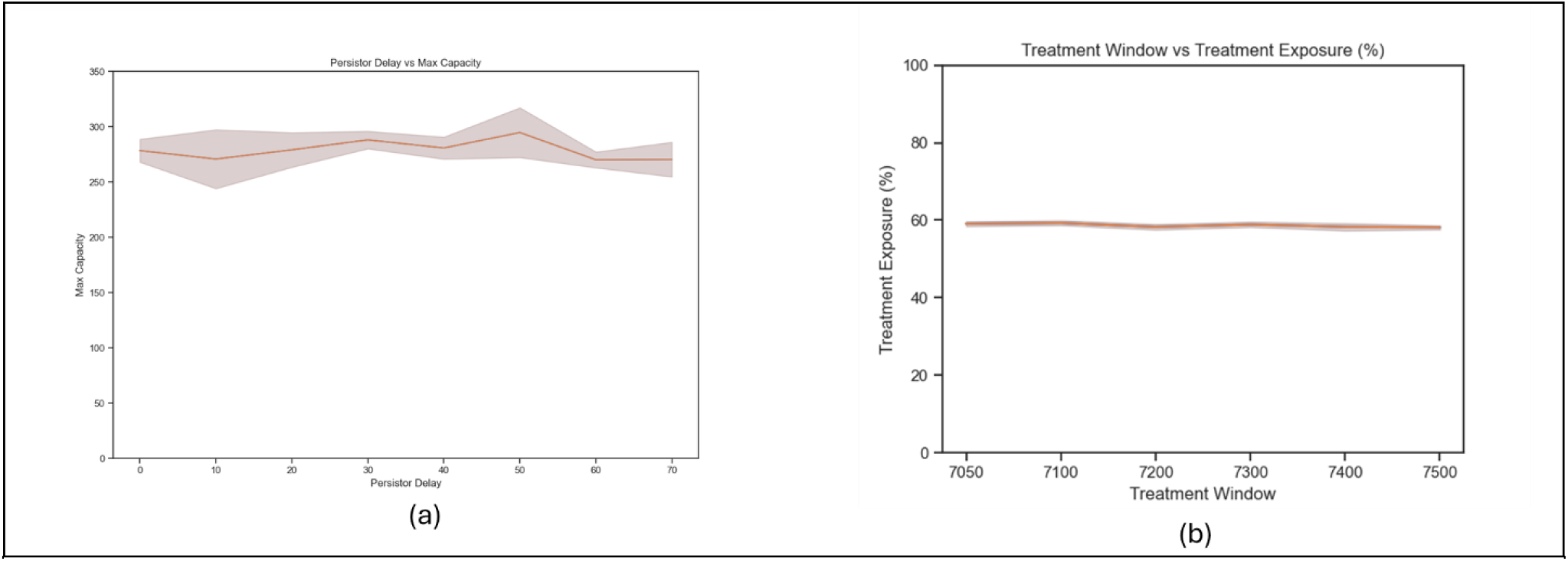
(a) The left plot presents the effect of delay in the onset of epigenetic resistance. In this model, it has no effect. (b) Provides evidence that the net treatment exposure is constant.

**Supplementary Figure 4.**
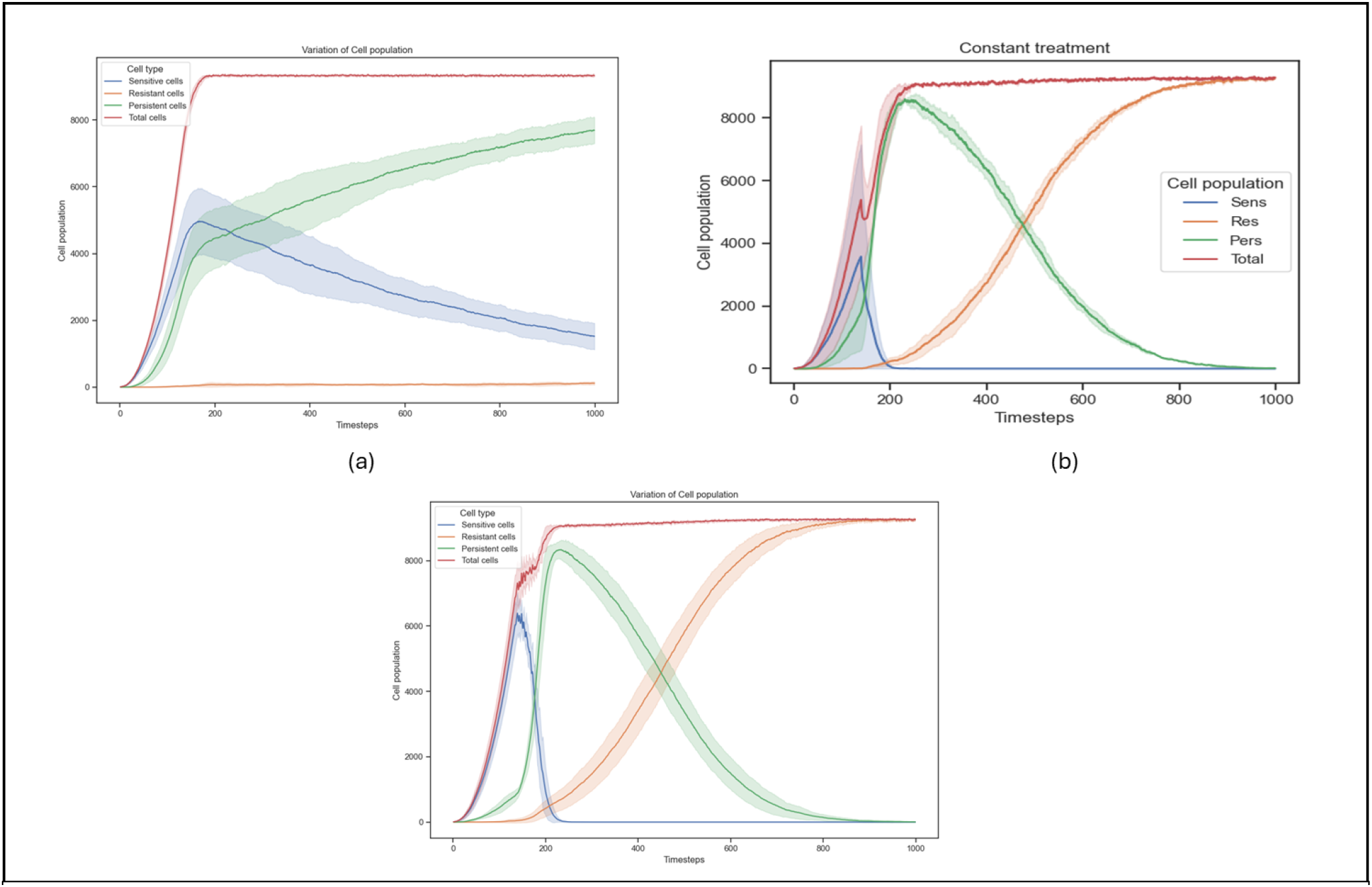
(a) The left plot presents a comparison between the treatment window and the duration until the tumor progresses.(b) The middle plot illustrates the marginal increments in both the treatment window and the time until the tumor reaches the progressive stage. (c) The right plot contrasts the treatment window with the number of treatment cycles. This comprehensive visual representation aids in understanding the interplay between treatment timelines and tumor progression.

Plasticity in cells is crucial for maintaining fitness under environmental stress. For this model, the plasticity of tumor cells is represented by epigenetic resistance, where a subset of cells is decreased in their susceptibility through downregulating their metabolism. When treatment is initiated, there is a delay in the development of persister phenotypes. Epigenetic modifications can be used to regulate and modulate the activity of the tumor, as they can create a lag phase in the development of complete resistance. However, simulations revealed that, despite very dramatic increases in the lag to emergence of epigenetic resistance, overall time for tumor to achieve maximum growth potential was unaffected—a surprising result that implies a lack of direct impact on the tumor’s overall growth rate. Figure (b) indicates that the percentage of time the tumor is exposedto treatment remains that same and hence does not skew results.

### Interplay of Carrying Capacity and Resistance Development

The simulation employs a 100 × 100 lattice, accommodating up to 10,000 cells, to model stochastic interactions among heterogeneous cell populations. This model adheres to dynamic population interaction principles, incorporating resource limitations such as nutrients and oxygen. As the population approaches the carrying capacity, stabilizing at approximately 9,000 cells, the tumor’s heterogeneity evolves, demonstrating increased resistance to applied therapeutic interventions. This progression towards resistance occurs irrespective of the treatment modality, albeit at varying rates. Constant, maximal treatment accelerates resistance development, while adaptive and intermittent therapies decelerate this process.

### Relationship of Tumor Size and Treatment Efficacy

Empirical evidence from multiple simulations with varying tumor sizes indicates that adaptive therapies exhibit enhanced efficacy when applied to larger tumors. The PFS positively correlates with tumor size. This relationship is further corroborated by an increase in the number of treatment cycles as tumor size increases. A plausible explanation for this phenomenon is the higher proportion of treatment-sensitive cells in larger tumors, rendering a greater percentage of the tumor susceptible to therapeutic intervention.

**Supplementary Figure 5.**
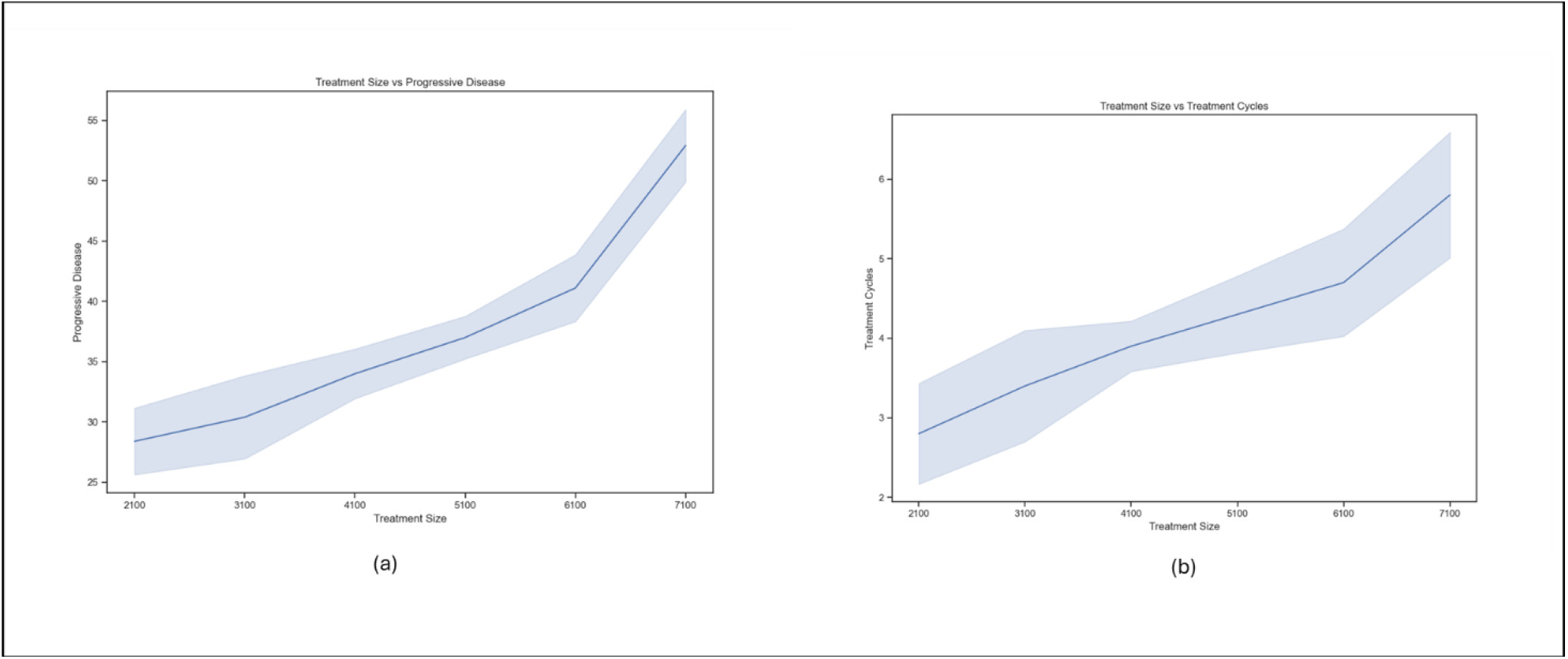
(a) A comparison of the treatment size and the duration until the tumor progresses. (b) A comparison between the treatment size with the number of treatment cycles.

### Treatment Window Optimization

**Supplementary Figure 6.**
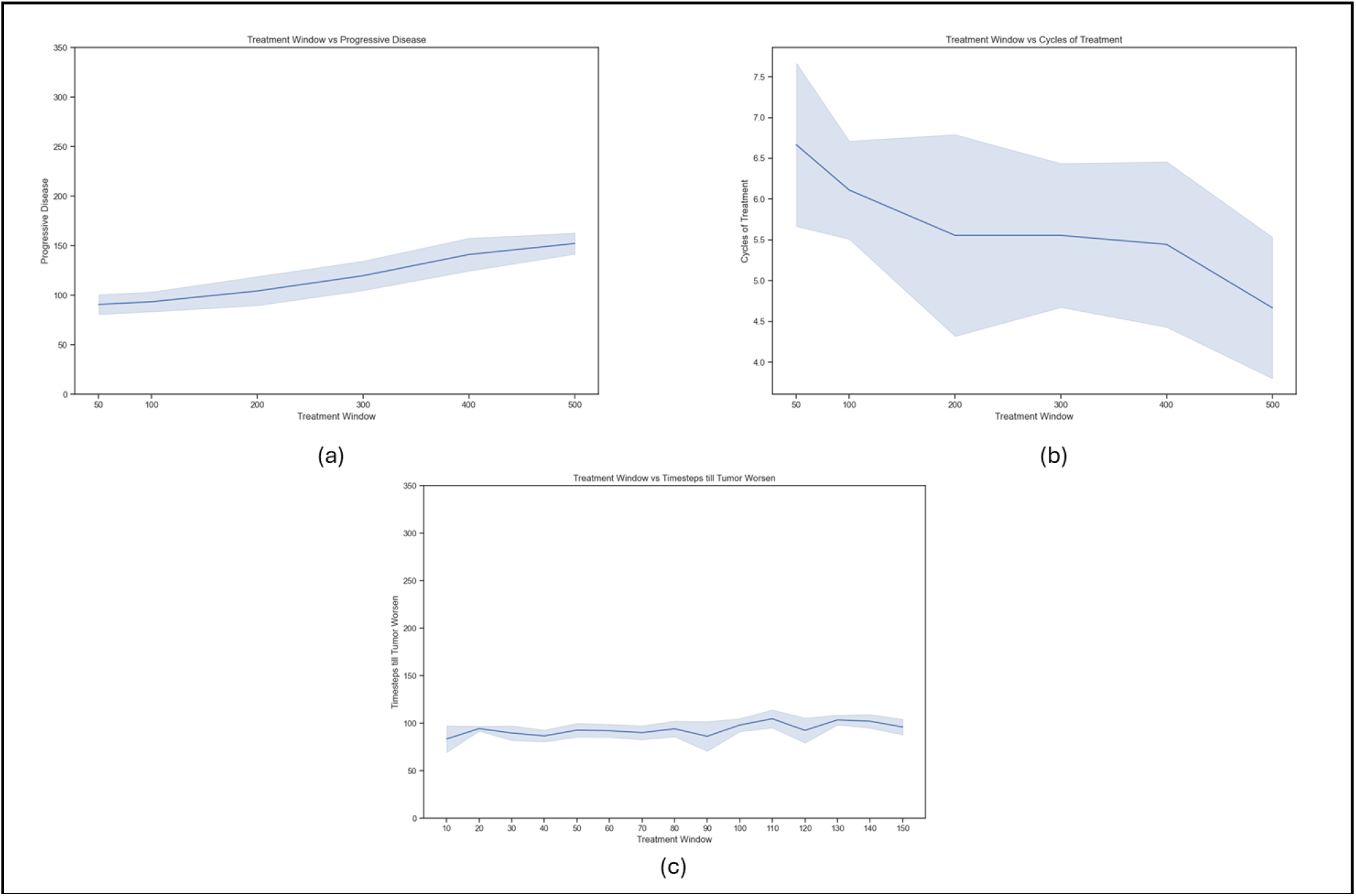
(a) The left plot presents a comparison between the treatment window and the duration until the tumor progresses. (b) The middle plot illustrates the marginal increments in both the treatment window and the time until the tumor reaches the progressive stage. (c) The right plot contrasts the treatment window with the number of treatment cycles. This comprehensive visual representation aids in understanding the interplay between treatment timelines and tumor

The treatment window, a crucial parameter in adaptive therapy, defines the range of cell populations within which treatment is administered. This window plays a pivotal role in maintaining the equilibrium of cell populations and optimizing treatment efficacy. Experimental results demonstrate that excessively wide treatment windows can lead to an unfavorable reduction in the proportion of sensitive cells, potentially compromising the objective of prolonging tumor stabilization. Conversely, overly narrow windows fail to significantly extend the time to tumor progression. Analysis of the data reveals a non-linear relationship between treatment window size and tumor progression. Marginal increases in the treatment window yield diminishing returns in terms of delaying tumor progression. Notably, substantial expansions of the treatment window result in only modest increases in time to progression. Furthermore, an inverse relationship is observed between the treatment windowsize and the number of treatment cycles required, suggesting that comparable therapeutic outcomes can be achieved with less frequent interventions when utilizing larger treatment windows. These findings underscore the importance of judicious selection of treatment parameters in adaptive therapy protocols, balancing the trade-offs between treatment frequency, tumor response, and the development of resistance.

## Notes

https://github.com/A-Binoy/Cancer-sim-python

### Competing Interest Statement

The authors have declared no competing interest.

### Funding

- National Cancer Institute: U54 CA274507

## References

[1] A. R. A. Anderson and V. Quaranta, “Integrative mathematical oncology,” Nat Rev Cancer, vol. 8, no. 3, pp. 227–234, Mar. 2008, doi: 10.1038/nrc2329.

[2] R. C. Rockne and J. G. Scott, “Introduction to Mathematical Oncology,” JCO Clin Cancer Inform, no. 3, pp. 1–4, Dec. 2019, doi: 10.1200/CCI.19.00010.

[3] S. Wang, J. Lei, X. Zou, and S. Jin, “Integrating multiscale mathematical modeling and multidimensional data reveals the effects of epigenetic instability on acquired drug resistance in cancer,” 2025, doi: 10.1371/journal.pcbi.

[4] A. Araujo et al., “Quantification and optimization of standard-of-care therapy to delay the emergence of resistant bone metastatic prostate cancer,” Cancers (Basel), vol. 13, no. 4, pp. 1–16, Feb. 2021, doi: 10.3390/cancers13040677.

[5] J. L. Gevertz, J. M. Greene, S. Prosperi, N. Comandante-Lou, and E. D. Sontag, “Understanding therapeutic tolerance through a mathematical model of drug-induced resistance,” NPJ Syst Biol Appl, vol. 11, no. 1, p. 30, Apr. 2025, doi: 10.1038/s41540-025-00511-3.

[6] R. A. Gatenby, A. S. Silva, R. J. Gillies, and B. R. Frieden, “Adaptive Therapy,” Cancer Res, vol. 69, no. 11, pp. 4894–4903, Jun. 2009, doi: 10.1158/0008-5472.CAN-08-3658.

[7] S. V. Sharma et al., “A Chromatin-Mediated Reversible Drug-Tolerant State in Cancer Cell Subpopulations,” Cell, vol. 141, no. 1, pp. 69–80, 2010, doi: 10.1016/j.cell.2010.02.027.

[8] M. Ramirez et al., “Diverse drug-resistance mechanisms can emerge from drugtolerant cancer persister cells,” Nat Commun, vol. 7, Feb. 2016, doi: 10.1038/ncomms10690.

[9] K. Kochanowski, L. Morinishi, S. J. Altschuler, and L. F. Wu, “Drug persistence – From antibiotics to cancer therapies,” Aug. 01, 2018, Elsevier Ltd. doi: 10.1016/j.coisb.2018.03.003.

[10] X. X. Sun and Q. Yu, “Intra-tumor heterogeneity of cancer cells and its implications for cancer treatment,” Oct. 01, 2015, Nature Publishing Group. doi: 10.1038/aps.2015.92.

[11] B. Zhitomirsky et al., “Abstract 2859: High-resolution lineage tracing for the study of cancer drug persistence at the single-cell level,” Cancer Res, vol. 83, no. 7_Supplement, pp. 2859–2859, Apr. 2023, doi: 10.1158/1538-7445.AM2023-2859.

[12] P. J. Tebon et al., “Drug screening at single-organoid resolution via bioprinting and interferometry,” Nat Commun, vol. 14, no. 1, Dec. 2023, doi: 10.1038/s41467-023-38832-8.

[13] D. Huang et al., “High-Speed Live-Cell Interferometry: A New Method for Quantifying Tumor Drug Resistance and Heterogeneity,” Anal Chem, vol. 90, no. 5, pp. 3299– 3306, Mar. 2018, doi: 10.1021/acs.analchem.7b04828.

[14] E. A. Oliveira et al., “Epigenetic Heritability of Cell Plasticity Drives Cancer Drug Resistance through a One-to-Many Genotype-to-Phenotype Paradigm Running title: Epigenetic heritability of cell plasticity drives resistance Conflict of Interest”, doi: 10.1158/0008-5472.CAN-25-0999/3621208/can-25-0999.pdf.

